# Breeding and hibernation of captive meadow jumping mice (*Zapus hudsonius*)

**DOI:** 10.1101/2020.10.02.323386

**Authors:** Ethan A. Brem, Alyssa D. McNulty, William J. Israelsen

## Abstract

Hibernating mammals exhibit unique metabolic and physiological phenotypes that have potential applications in medicine or spaceflight, yet our understanding of the genetic basis and molecular mechanisms of hibernation is limited. The meadow jumping mouse, a small North American hibernator, exhibits traits – including a short generation time – that would facilitate genetic approaches to hibernation research. Here we report the collection, captive breeding, and laboratory hibernation of meadow jumping mice. Captive breeders in our colony produced a statistically significant excess of male offspring and a large number of all-male and all-female litters. We confirmed that short photoperiod induced pre-hibernation fattening, and cold ambient temperature facilitated entry into hibernation. During pre-hibernation fattening, food consumption exhibited non-linear dependence on both body mass and temperature, such that food consumption was greatest in the heaviest animals at the coldest temperatures. Meadow jumping mice exhibited a strong circadian rhythm of nightly activity that was disrupted during the hibernation interval. We quantified the length and timing of torpor bouts and arousals obtained from an uninterrupted recording of a hibernating female. Over a 90.6 day hibernation interval, torpor bouts ranged from 2.1 to 12.8 days (mean 7.7 days), and arousal length was relatively constant with a mean length of 9.6 hours. We conclude that it is possible to study hibernation phenotypes using captive-bred meadow jumping mice in a laboratory setting.

## Introduction

Daily torpor and hibernation form a continuum of phenotypes that is widespread throughout many orders of mammals [1, 2], suggesting that heterothermy is an ancestral trait that was maintained during the evolution of endothermy [3]. Mammalian hibernators include species of bears, bats, rodents, marsupials, and even primates [4]. Although the state of torpor employed by hibernating animals – profoundly reduced body temperature and significantly slowed metabolic rate – has attracted attention for its potential therapeutic applications in human medicine and spaceflight [5, 6], little is known about its underlying molecular mechanisms or genetics [1]. Hibernation research has historically been closely aligned with ecology and ecological physiology, with the organisms used in hibernation research selected from wild populations locally available to investigators. Traditional species of interest – including bears, bats, ground squirrels, and dormice – exhibit a prolonged hibernation interval fueled by accumulated fat stores, but such species have long generation times and are therefore not readily amenable for genetic studies of hibernation phenotypes. The difficulty in investigating the genetic basis of hibernation is illustrated by the fact that, to our knowledge, only one such study has been published, which identified candidate genes affecting the timing of hibernation onset in the thirteen lined ground squirrel [7]. The golden (or Syrian) hamster (*Mesocricetus auratus*) is more amenable to genetic approaches [8, 9], but this and other hamster species hoard food and hibernate in a lean state with relatively shorter bouts of torpor [10–14], which would not allow us to study our phenotypes of interest.

As part of a long-term goal to accelerate genetic and molecular aspects of hibernation research, we selected the meadow jumping mouse (*Zapus hudsonius*) as a model organism. The meadow jumping mouse is a small hibernator native to North America; it exhibits pre-hibernation fat storage and a deep hibernation phenotype with long torpor bouts and short interbout arousals [15–18]. The meadow jumping mouse produces multiple litters of offspring per year and is known to reproduce prior to its first hibernation [19, 20], potentially opening the door to genetic experimental approaches that involve breeding and thus require short generation times. Because the meadow jumping mouse uses modifiable environmental cues such as day length and temperature to determine the timing of hibernation [21], this species also enables convenient laboratory study of hibernation phenotypes. Despite the potential utility of the meadow jumping mouse as a model hibernator we were unaware of any reported systematic breeding of this species. Here we report our successful efforts to collect wild meadow jumping mice and establish a captive breeding colony. We investigated the conditions required for induction of fat accumulation and hibernation in the laboratory. Both short photoperiod and low temperature contributed to the weight gain and successful hibernation of meadow jumping mice, which confirmed previous work [21]. We measured food consumption during pre-hibernation fattening and hibernation and identified an interaction effect between animal body mass and housing temperature, in that cold temperature increases pre-hibernation food consumption to the greatest degree in the heaviest mice. We determined the amount and timing of daily locomotor activity before and during hibernation, and quantified the lengths of torpor bouts and the lengths and timing of interbout arousals from the complete hibernation interval of a captive animal. Torpor bout length changed throughout hibernation, but arousal length remained constant and the timing of arousals with respect to photoperiod showed an unexpected degree of regularity.

## Materials and Methods

### Animal collection and health testing

Animal work described in this manuscript has been approved and conducted under the oversight of the UT Southwestern Institutional Animal Care and Use Committee (protocol 2014-152 & 2015–101240) and the Massachusetts Institute of Technology Committee for Animal Care (protocol 0514-048-17 & 0718-064-21). Animal collection was approved by the Massachusetts Division of Fisheries and Wildlife and conducted according to the Guidelines of the American Society of Mammalogists for the use of wild mammals in research [22]. Wild meadow jumping mice were collected on the Bolton Flats Wildlife Management Area, near Bolton, MA, USA. Live traps (LFA Folding Trap, HB Sherman, Tallahassee, FL, USA) were baited with a mixture of peanut butter and rolled oats and left open overnight. Traps were checked near dawn and left closed during the day. Species other than meadow jumping mice were recorded and released. To mitigate risk of zoonotic illness, masks were worn in the field while working with occupied traps and while handling captive animals of unknown disease status and their soiled bedding, as informed by established guidelines [23]. Occupied traps were washed using a bleach solution prior before being returned to the field; this was a precaution against zoonotic disease and allowed collection of *Zapus* feces from clean traps for pathogen testing. Following capture, health screening was performed for each animal by MIT veterinary staff. Fresh feces from the trap were used for parasite screening (via fecal float test) and commercial pathogen screening, which included the Charles River Laboratory Mouse FELASA Complete PRIA panel with additional PCR tests for mites, Leptospira, and New World Hantavirus (Charles River Laboratories, Wilmington, MA, USA). In the first year, fecal samples were PCR tested by a second laboratory for Hantavirus (Assay #S0135, Zoologix, Chatsworth, CA, USA). Additional endo- and ectoparasite screening was performed via fur pluck and anal tape exam. All animals were treated with ivermectin for 4 weeks upon entry into the colony. Topical pyrethrin was used as an ectoparasiticide the first year of collection, and animals positive for Giardia parasite were successfully treated with a course of metronidazole in the third year of collection.

### Standard housing and diet

The breeding colony was housed at ~20° C and simulated summer photoperiod (16 hours light, 8 hours dark). Animals were housed individually in standard, filter-top plastic rodent cages (45 cm long × 24 cm wide × 16 cm high) with corncob bedding, a red plastic rodent shelter, and Enviro-dri bedding paper (Shepard Specialty Papers, Watertown, TN, USA). Cotton nestlets were not used due to instances of cotton fibers causing penile strangulation. Physical enrichments, including wood blocks, were sometimes included. The diet in current use is Teklad 2019 (Envigo, Indianapolis, IN, USA); water and rodent chow were provided ad libitum and the diet was enriched with small amounts of commercial animal enrichment foods such as whole sunflower seeds.

### Captive breeding

Breeding animals were housed in large plastic cages (55 cm long × 37.5 cm wide × 20.5 cm high). Each breeding cage contained two custom nest boxes (as described previously, but without instrumentation [24]), Enviro-dri bedding paper for each nest box, additional physical enrichment such as a red plastic shelter or wood block, and food and water. Typically, a potential breeder (male or female) was introduced into the breeding cage and provided at least one day to acclimate before the second animal was introduced. Following pairing, the male was removed after around 3 days with the female or at the first sign of incompatibility.

### Hibernation experiments

For hibernation experiments, animals were housed in environmental chambers (Rodent Incubator model RIS33SD, Powers Scientific, Pipersville, PA) set to the desired photoperiod and temperature using the integrated digital controls. When applicable, the absolute timing of the photoperiod relative to the rotation of the earth was not changed at the onset of US daylight saving time. One control group was housed in a standard animal room in the vivarium. Animals were housed in the same cages used for routine housing (45 cm long × 24 cm wide × 16 cm high), but they were provided with a nest box and Enviro-dri paper for use as a hibernaculum. Food and water were provided ad libitum for the duration of the hibernation experiments. The diet consisted of a standard rodent chow – Teklad 2018 (Envigo, Indianapolis, IN, USA) – plus two small sunflower seeds per day if the seeds from the previous day had been consumed. Visual observation of uneaten sunflower seeds signaled that the animal was likely torpid. Animals and food were weighed once or twice weekly (depending on experiment) near the end of the light phase using a digital laboratory scale to a resolution of 0.1 grams. Food was weighed using the same scale, and the average daily food consumption for each period (typically 3 or 4 days) was determined by dividing the mass of food consumed by the number of days between measurements. The daily ration of two sunflower seeds (excluding uneaten shells) weighed less than 0.1 g and was ignored in the food consumption calculations. Male mice were used for most experiments due to their excess numbers in in the colony and lesser relative importance in ongoing breeding efforts.

### Activity monitoring

Activity outside of the nest box was detected for each animal using a passive infrared (PIR) motion sensor mounted on the underside of the cage lid. Voltage outputs from the motion sensors were recorded in 5-minute increments using a Sable Systems UI-3 Universal Interface and ExpeData software (Sable Systems, North Las Vegas, NV, USA). Motion recordings were paused or stopped when cages were opened for weighing of food and animals to minimize spurious motion detection events. Due to software and computer memory limitations, recordings were saved and restarted weekly to minimize data loss. For purposes of comparison to food consumption and body mass data, each day of motion started at the onset of the dark phase and ended at the end of the light phase, when body mass data were obtained. Only those data available from complete or near-complete 24-hour recording periods were used.

Nest box temperature measurements were made at 30 second intervals using a K-type thermocouple and amplifier (Part Numbers 270 and 269, Adafruit Industries, New York City, NY, USA). The thermocouple probe was attached to the underside of the nest box lid, so that the probe was directly above the hibernating animal. The reference temperature was provided by the amplifier circuitry which was located inside the environmental chamber but outside of the cage. Data were recorded by a python script running on a single board computer (Raspberry Pi Model B+, Raspberry Pi Foundation, Cambridge, United Kingdom). The data were cleaned to remove bad data points and the difference between nest box and chamber temperature was plotted in R. We quantified bouts of torpor and arousals such that the arousal began when the nest box temperature differential first increased above baseline, and the arousal ended (and torpor began) when the temperature differential of an occupied nest box first began to decline. Due to high frequency noise in the data, the temperature inflection points were determined manually to the nearest 15 minutes.

### Data analysis and visualization

Data analyses and visualization were performed using Microsoft Excel (Microsoft Corporation, Redmond, WA, USA), GraphPad Prism version 8.4.3 (GraphPad Software, San Diego, CA, USA), and R version 3.6.0 [25] with the following packages: ggplot2 [26], lubridate [27], ggmap [28], and circular [29]. To estimate the probability of encountering the observed number of single-sex litters, 10 million sets of 32 litters of the same size distribution as those recorded in our colony were generated in R. For each litter, the sex of each pup was drawn from a binomial distribution using the rbinom() function, with the underlying probability of obtaining a male set at 0.5 (expected) or 0.6615385 (observed in our colony). The sex ratios of the simulated sets of litters were used to generate histograms, and the number of single-sex litters obtained in each of the 10 million sets of 32 litters was counted and used to determine the probability of obtaining the observed number of single-sex litters at each underlying sex ratio.

## Results

### Trapping and health assessment

The meadow jumping mouse is a member of superfamily Dipodoidea, which includes other jumping mice, birch mice, and jerboas. Meadow jumping mice are found throughout broad regions of North America, including much of the eastern United States, Canada, and Alaska (Fig 1A). Close North American relatives include other hibernating species in the same genus, the western jumping mouse (*Zapus princeps*) and Pacific jumping mouse (*Zapus trinotatus*), which both diverged from the meadow jumping mouse less than 3 million years ago [30]. The jumping mice are only distantly related to the gerbils and true mice and rats of superfamily Muroidea that are commonly used as laboratory models.

**Fig 1.**
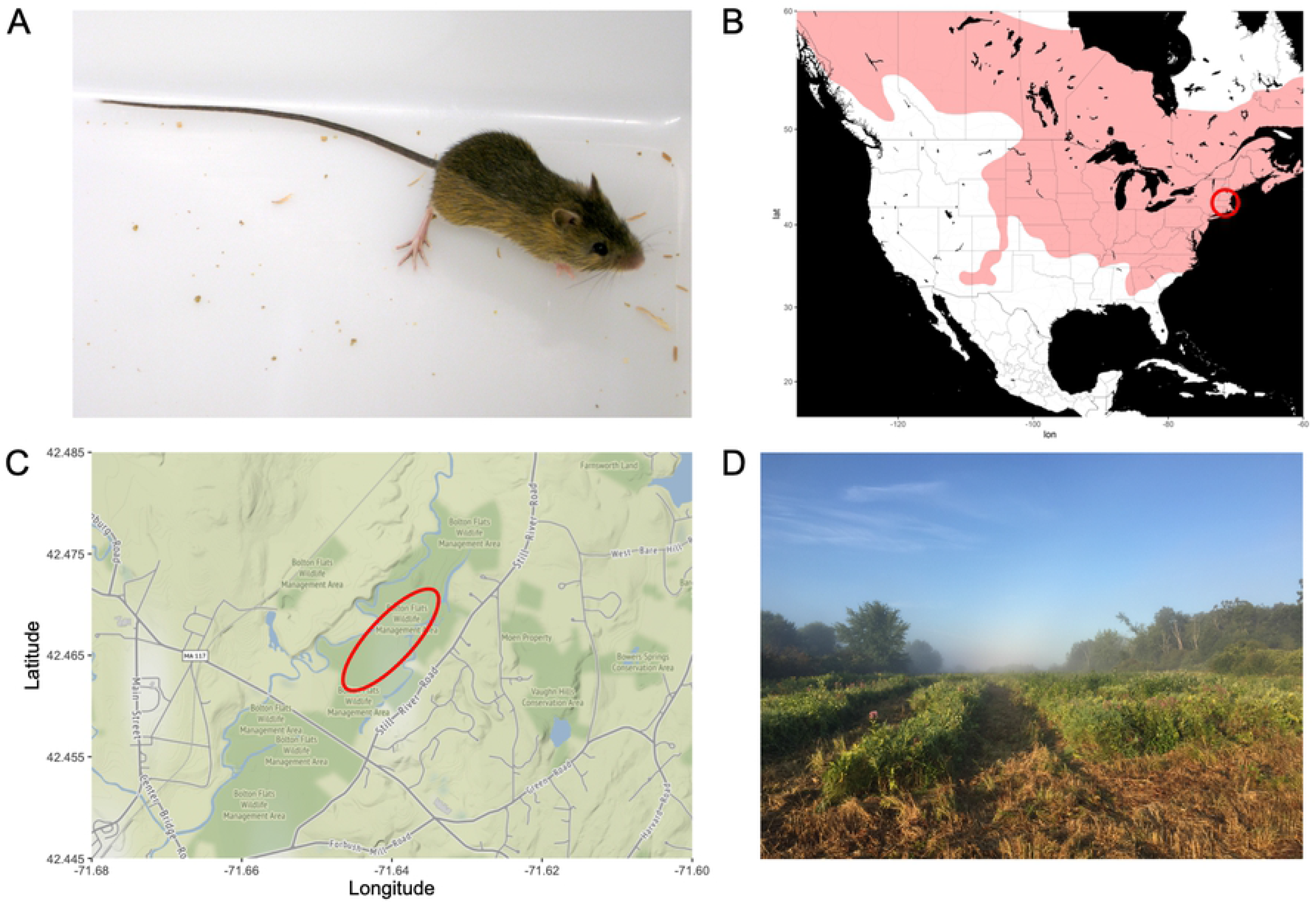
Appearance, range, and collection location of the animals used in this study. (**A**) An adult meadow jumping mouse, showing the long tail and large hind limbs used for jumping locomotion. (**B**) The range of *Zapus hudsonius* in much of North America. The collection location is marked with a red circle. (**C**) Bolton Flats Wildlife Management Area, with the trapping location indicated with a red ellipse. (**D**) Photograph showing typical late summer habitat in the trapping area, shown here after some of the meadow was mowed by the state wildlife agency.

In the Northeastern United States, meadow jumping mice can be found in grassy or sedge meadows near water and wetlands [15, 31, 32]. We identified a trapping locale with appropriate habitat on the Bolton Flats Wildlife Management Area near Bolton, Massachusetts, USA (Fig 1B). In Massachusetts, the active season during which *Zapus* can be collected generally spans from May or June through the end of summer or early autumn [32]. Meadow jumping mice were collected in the late summer (August-September) of three seasons using Sherman live traps. Our trapping efforts resulted in a capture rate of 1.47 *Zapus* per 100 trap-nights over three years, and complete trapping results for all species captured are presented in Table 1.

**Table 1.** Species captured during three years of *Zapus* collection.

| Genus | Common Name | 2014 | 2015 | 2019 | Total |
| --- | --- | --- | --- | --- | --- |
| *Zapus* | Meadow Jumping Mouse | 15 | 13 | 25 | 53 |
| *Sorex* | Shrew | 37 | 3 | 17 | 57 |
| *Peromyscus* | White-footed or Deer Mouse | 173 | 9 | 14 | 196 |
| *Blarina* | Short-tailed Shrew | -- | -- | 8 | -- |
| *Microtus* | Meadow Vole | -- | -- | 2 | -- |
| *Blarina* + *Microtus*<sup>a</sup> | Short-tailed Shrew + Vole<sup>a</sup> | 30 | 11 | 10 | 51 |
| *Mustela* | Weasel | 1 | 1 | 0 | 2 |
| --<sup>b</sup> | Frog<sup>b</sup> | 1 | 2 | 67 | 70 |
| **Summary** |  |  |  |  |  |
| Trap Nights |  | 977 | 684 | 1938 | 3599 |
| Total Captures |  | 257 | 39 | 133 | 429 |
| Total Captures Per 100 Trap Nights |  | 26.3 | 5.7 | 6.9 | 11.9 |
| *Zapus* Captures per 100 Trap Nights |  | 1.5 | 1.9 | 1.3 | 1.5 |

All captures occurring during three seasons collection on the Bolton Flats Wildlife Management Area, Massachusetts, United States of America. All species other than *Zapus* were released.
*Peromyscus* captures were reduced following historic snowfall during the winter of 2014-15.
Significant flooding in 2019 resulted in many frog captures but did not impair *Zapus* collection.
<sup>a</sup>We did not differentiate between these two species during the first two collection years.
<sup>b</sup>We did not attempt to identify the frogs.

Meadow jumping mice have a low parasite load [15], and *Zapus* species are not known to be vectors for zoonotic diseases including Hantavirus [33, 34]. However, standard precautions were taken during field work and the initial stages of animal husbandry because *Peromyscus* spp. in the trapping area are known reservoirs of Hantavirus [35] and the potential for the presence of additional pathogens in the meadow jumping mice was unknown. Once collected, the captured animals were quarantined and tested for zoonotic diseases, an extensive panel of rodent pathogens, and endo- and ectoparasites. The captive animals were also prophylactically treated with the anti-parasitic ivermectin and, in some cases, topical pyrethrin as previously reported [36]. Upon entry, a few animals tested positive for a limited number of rodent pathogens and parasites including Mouse Adenovirus, *Helicobacter ganmani*, *Klebsiella oxytoca*, *Giardia*, coccidian enteroparasites, and a non-typical species of mites (S1 Table); the mites were probably *Dermacarus hypudaei* in its non-feeding form, as reported previously [15]. The few animals that initially tested positive for Mouse Adenovirus tested negative upon repeated testing and no pathogens or parasites were passed to via bedding to sentinel animals (*Mus musculus*) used in routine facility pathogen monitoring. Animals positive for *Giradia* parasites were either excluded from the colony or, in the final season, successfully treated with metronidazole. All captured animals tested negative for zoonotic pathogens including Hantavirus, cytomegalovirus (CMV), and *Leptospira*.

### Housing and breeding

Meadow jumping mice are solitary in the wild except when raising young [15], and they have been housed individually in rodent cages by a previous investigator [21]. We have successfully housed individual meadow jumping mice in large plastic rodent cages (45 cm long × 24 cm wide × 16 cm high) that provide more space for saltatory locomotion than standard laboratory mouse cages (Fig 2A). Custom-built plastic, hinged-top nest boxes (Fig 2B), previously reported by us and similar in design to those used for jerboas [24, 37], provide a burrow-like refuge during breeding and hibernation. Standard rodent shelters and paper bedding strips, which mimic the grass often used for nest building in the wild [19, 38], are provided during routine housing. Wild meadow jumping mice subsist on a diet of grass seed and other seeds, which are heavily supplemented by insects, berries, and the terrestrial fungus *Endogone* [19, 39, 40]. The laboratory diet employed by us consists of water and a standard laboratory rodent chow with a limited amount of sunflower seeds and/or other commercially available lab animal diet enrichment foods. Given the leaping ability of the jumping mice, we have found it convenient to work with them by placing their cages into large plastic bins (60 cm long × 40 cm wide × 37 cm high) so that the animal is contained if it jumps out of the cage (Fig 2C). A low-stress method for weighing or transferring animals between cages is to have them hop into a plastic beaker as a transfer device (Fig 2D-E). The animals can be handled and restrained in the fashion usually employed for laboratory mice or other small rodents such as *Peromyscus* [41], but we note that meadow jumping mice attempt to jump and kick powerfully if lifted by the tail. Such handling does not break off part of the tail as with the California pocket mouse [42], but we have found it best to grasp by the base of the tail and minimize unsupported lifting of the hind end if this method of handling is necessary.

**Fig 2.**
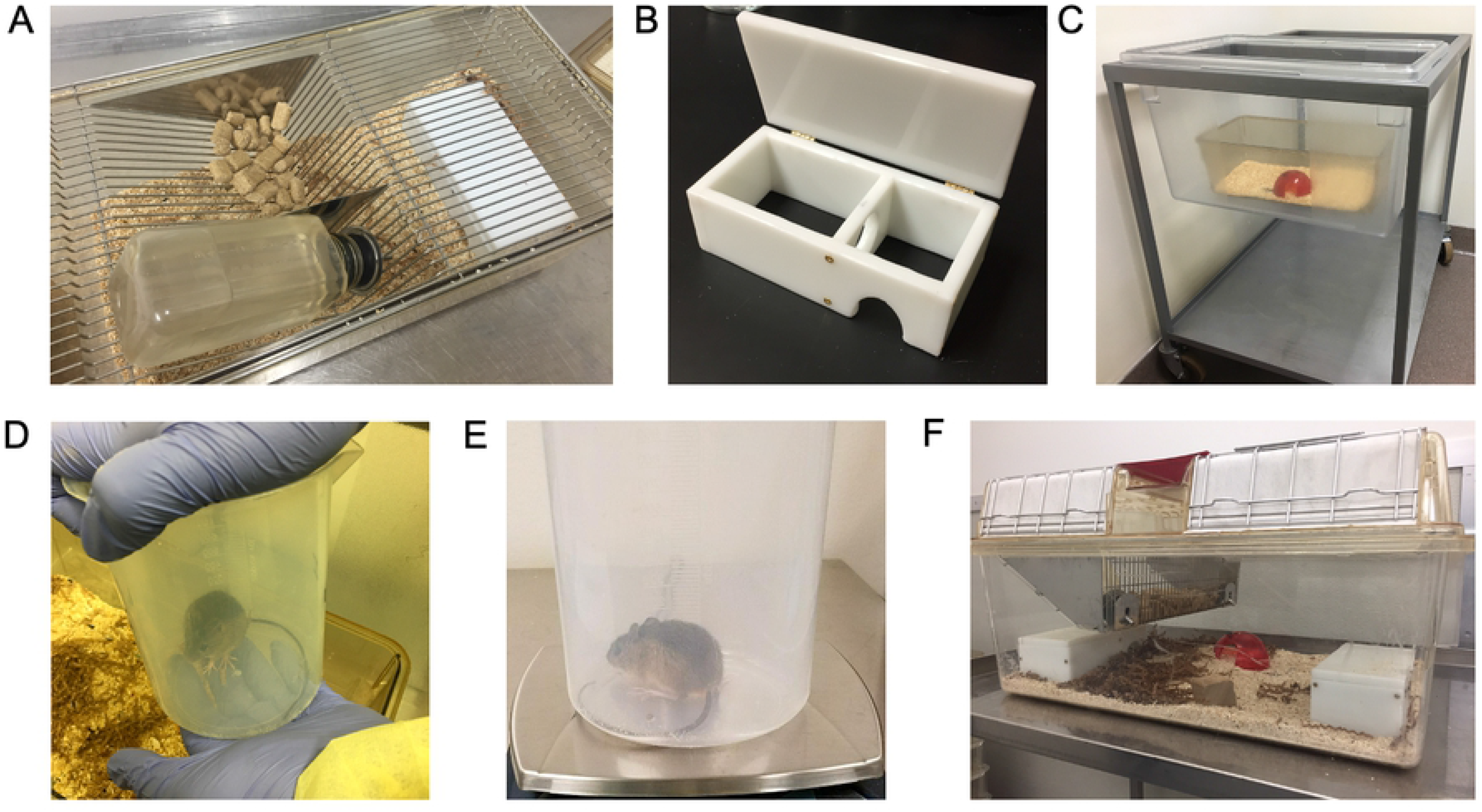
Laboratory housing and handling of the meadow jumping mouse. (**A**) A cage used for single housing of *Zapus*; the white nest box fits inside this cage for hibernation experiments. (Cage lid not shown). (**B**) Interior of nest box, showing entrance, hinged lid, and two interior chambers. (**C**) Placing the cage inside a large plastic bin provides containment during animal handling. This demonstration cage contains the type of red plastic shelter provided during routine housing. (**D**) We use a plastic beaker for low-stress animal handling when restraint is not required. (**E**) Body mass can be conveniently determined with the animal contained inside a plastic beaker. (**F**) The large cages used for breeding contain two nest boxes and additional enrichment.

We house our breeding animals in large plastic cages (55 cm long × 37.5 cm wide × 20.5 cm high) of the size typically used for guinea pigs at our institution; the animals are provided with two nest boxes, bedding material, and additional enrichment items (Fig 2F). These large cages allow space for interactions between the paired male and female and for the subsequent litter. Because meadow jumping mice do not pair bond, we house the male with the female for generally no more than 3 days. Failure to remove the male from the cage in a timely fashion invariably results in maternal cannibalism of any litter produced. The female is also very sensitive to human disturbance during the perinatal and postpartum period, which can also result in destruction of the litter. Maternal cannibalism in captivity and the tendency of the female to abandon the litter following disruption of the nest in the field have been previously reported [38, 39]. Despite these challenges, we found that meadow jumping mice breed successfully in captivity with appropriate care (Fig 3A).

**Fig 3.**
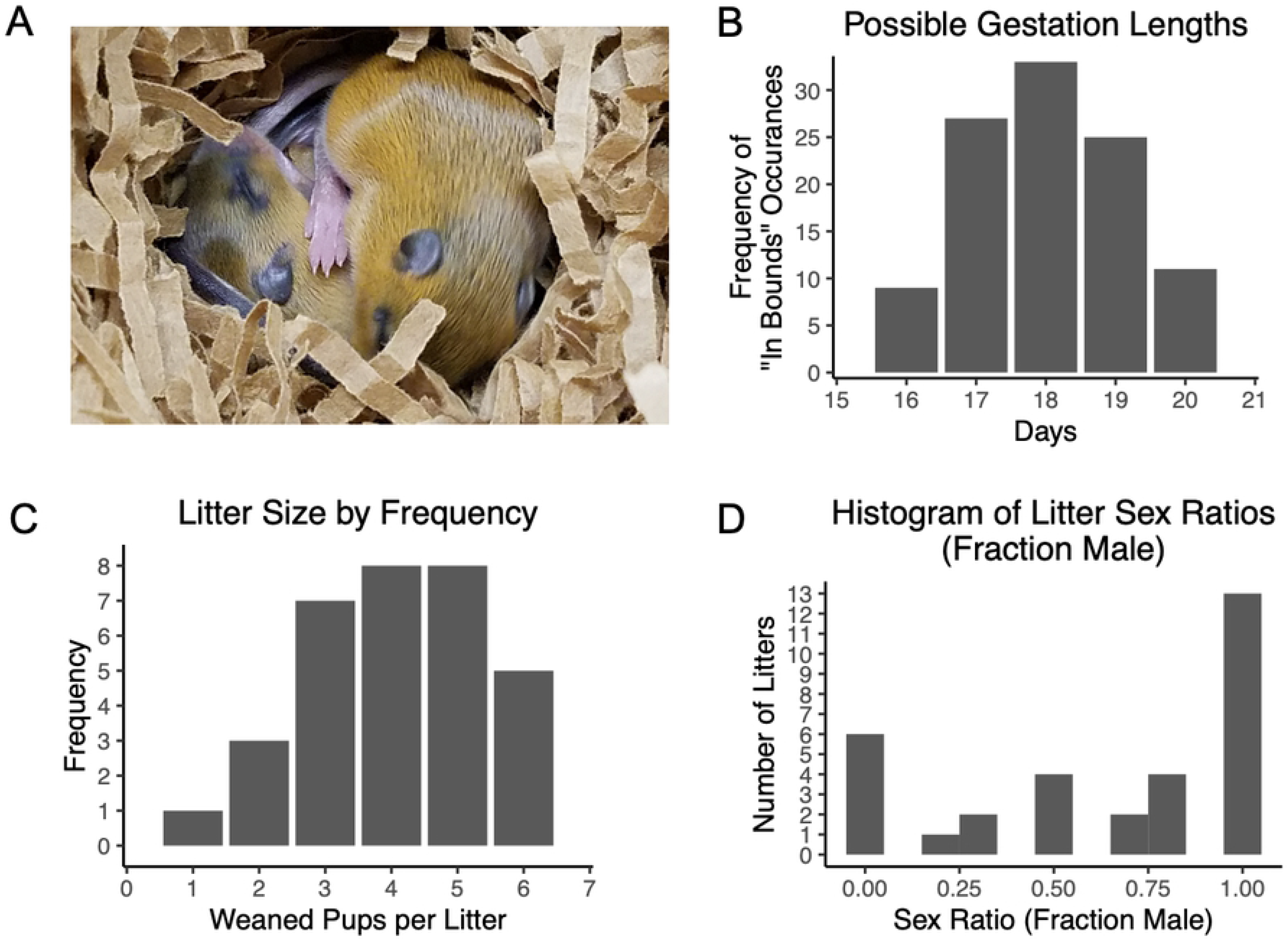
Captive breeding of *Zapus* results in a male-biased sex ratio and a high proportion of single-sex litters. (**A**) Meadow jumping mouse pups visible in an opened nest box. These animals are approximately 17 days old. (**B**) Histogram of possible gestation lengths from 39 successful matings; impossible lengths were removed from the frequency distribution according to the number of times they were out of bounds given the breeder pairing and separation dates. (**C**) The distribution of litter size (number of weaned pups) from each of 32 litters (mean ; 4 pups). (**D**) Histogram of litter sex ratio (fraction male) in our colony, determined from sexes of weaned animals in 32 litters. Note the unusually large number of all-female and all-male litters.

In the wild, meadow jumping mice become reproductively active upon emergence from hibernation and continue breeding throughout the summer. Although reports differ somewhat by location, meadow jumping mice populations exhibit up to two or three peaks of reproductive activity during each breeding season which may extend from May to September [19, 20, 32, 39, 43]. The litter sizes found in field studies ranged from 3 to 7 or 4 to 7 pups, with means of 4.5 or 5.7 pups per litter, depending on location [15]. Reported gestation lengths from 4 captured females were in the range of 17 to 21 days, with the longest gestation of 20 to 21 days possibly having been prolonged because that animal was lactating [19]. The relatively short gestation time allows females of this species to produce more than one litter per year, and females have been observed to breed during the same summer in which they are born [19, 20], indicating that hibernation is not required for females to reach sexual maturity.

Our captive breeding efforts provided additional data confirming the reported gestation lengths and also revealed a male-skewed sex ratio and a propensity for production of same sex litters. The pairing and separation dates from 39 successful breeding attempts showed that the most common possible gestation length was 18 days (Figs 3B and S1A), which falls within the previously reported range [19]. To avoid reducing the probability of breeding success, we did not investigate the presence or absence of a copulatory plug in this species by handling the paired females; however, we do not expect that the male produces a copulatory plug, as the male lacks some accessory glands [44] and evidence of mixed-paternity litters has been found in the closely-related Pacific jumping mouse, *Zapus trinotatus* [45]. Direct observation of copulation, or the identification of a copulatory plug that can be used to determine the time of mating, will be necessary to make a more direct determination of gestation length. During our breeding efforts, we recorded the sexes of 130 pups weaned from a total of 32 surviving litters. Of these, 86 were male and 44 were female, resulting in a sex ratio (fraction male) of 0.662. The observed sex ratio was significantly different from an expected sex ratio of 0.5 (p=0.0002895, exact binomial test), indicating a skew toward male offspring in our breeding conditions. The number of pups weaned per litter ranged from 1 to 6, with a mean litter size of 4 (Fig 3C). The maximum number of neonates observed was 8, which did not exceed the number of eight mammary glands present in this species. Surprisingly, of the 32 litters, 6 were all-female and 13 were all-male, providing a distribution of sex ratios per litter (Fig 3D) unlike the distributions that would be expected if pups were produced at the expected (0.5) or observed (0.662) sex ratios (S1B-C Fig). The total number of single-sex litters observed (19 of 32 litters) was significantly greater than would be expected from litters with pups produced at the expected (0.5) or observed sex ratios (0.662) (p < 1 × 10^−7^ and p = 8.2 × 10^−6^, respectively; S1D-E Fig). The size distribution of the single-sex litters is similar to that of all litters, with a mean of 4 and a maximum of 6 pups of the same sex (S1F Fig), suggesting that the unusually large proportion of single-sex litters is not simply an artifact of very small litter sizes. While significantly skewed sex ratios occur among many mammal species [46], the propensity of our meadow jumping mice to produce a high proportion of all-male and all-female litters is unexplained.

### Hibernation

#### Photoperiod controls hibernation phenotypes

The timing of breeding and hibernation in the meadow jumping mouse appears to be driven largely by day length [21, 47], which we suggest allows animals born early or late in the summer to prepare for hibernation at the appropriate time of year. The long photoperiod during summer inhibits pre-hibernation fattening in meadow jumping mice [21], and the animals remain in breeding condition under these conditions. Exposure to short days triggers the cessation of reproduction and promotes pre-hibernation weight gain [21].

We sought to further understand the effects of photoperiod and air temperature on initiation of hibernation in wild caught and captive-reared meadow jumping mice. Based on previous work, we expected that meadow jumping mice would not prepare for hibernation when housed under conditions approximating mid-summer day length at the latitude of the capture site. Accordingly, we found that animals born in the colony and housed at ~20° C and long photoperiod (16 hours light, 8 hours dark) remained at a summer weight and did not prepare for hibernation (Fig 4A). When plotted individually, these animals exhibited little variation in body mass over time (S2A Fig). Similarly, animals born to a female that was pregnant when captured in late summer did not fatten when housed at ~20° C and 16 hour photoperiod (Fig 4B, red lines), consistent with the observation that simulated summer conditions inhibit preparation for hibernation. In contrast, animals captured in August or September spontaneously fattened in captivity and maintained an increased body mass for several months (Fig 4B-C and S2B-C Fig), even when held under our standard simulated summers conditions (~20° C; 16 hours light, 8 hours dark). Some of the captured animals that had thus fattened in preparation for hibernation (Fig 4D) spontaneously entered torpor as indicated by adoption of the stereotypical curled posture (Fig 4E), external body temperature near ambient temperature of ~20 °C, and initial lack of responsiveness to external stimuli (Fig 4F). These observations imply that the environmental inputs experienced by the animals in the field prior to capture in August or September were sufficient to launch a putative physiological ‘hibernation program’ that ran to completion before the animals returned to summer body weight. Following a return to summer body weight, a second exposure to a shorter day length (12 hours light, 12 hours dark) at colony housing temperature (~20° C) was sufficient to again induce pre-hibernation fattening (Fig 4C). We note that the 12-hour light phase approximates astronomical conditions around the autumnal equinox (late September) in our capture area, which falls near the end of the meadow jumping mouse active season.

**Fig 4.**
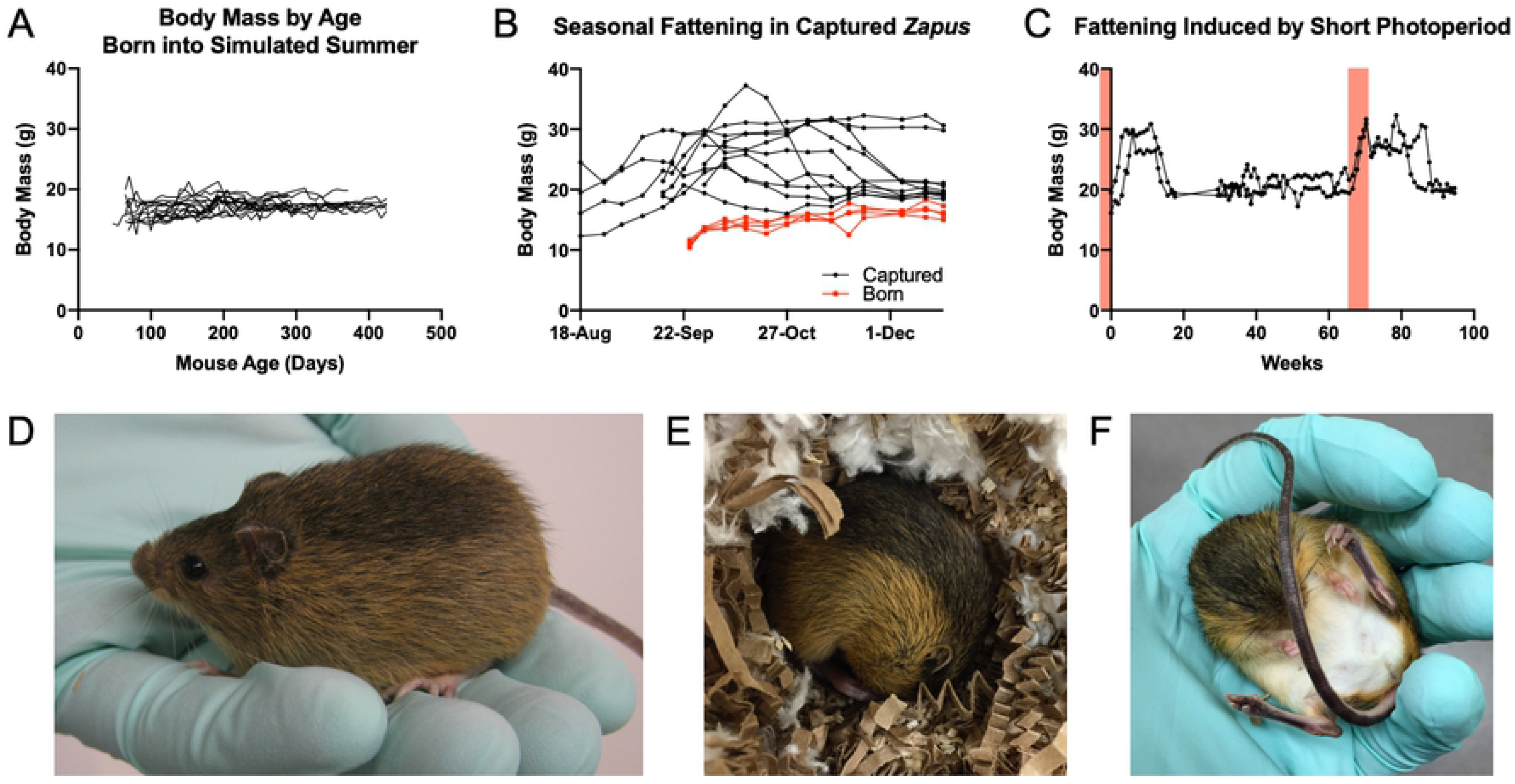
Photoperiod controls pre-hibernation fattening in wild-caught and captive-reared meadow jumping mice. (**A**) Body mass by age of 15 captive-born meadow jumping mice that were housed under long photoperiod (16L/8D) which inhibited pre-hibernation fattening. (**B**) Body mass of animals captured near the end of the active season (black lines) or born to a female that was pregnant while captured (red lines) and held at long photoperiod (16L/8D). Animals captured at the end of summer fattened for hibernation despite the long photoperiod; captive-born animals did not fatten. (**C**) Periods of exposure to short photoperiod (12L/12D, shaded red) are sufficient to induce pre-hibernation fattening in animals were held at long (16L/8D) photoperiod after capture. (**D**) A meadow jumping mouse that has prepared for hibernation by fattening. (**E**) A meadow jumping mouse curled up in typical hibernation posture. The nest box lid and nest were opened to view the animal. (**F**) A partially uncurled animal that was found torpid at 20° C ambient temperature. Full arousal takes around 10 minutes at this temperature [24].

#### Temperature affects pre-hibernation body mass and food consumption

We next investigated the importance of ambient temperature on hibernation induction in our captive-reared meadow jumping mice. Because a single photoperiod was to be used for the duration of this experiment, we selected a simulated mid-winter photoperiod (8 hours light, 16 hours dark) that we reasoned would be sufficient to both induce and maintain hibernation in our animals. Groups of five animals were housed under control conditions in an environmental chamber or standard housing room (20° C, 16 hours light, 8 hours dark), or under three induction conditions combining short photoperiod (8 hours light, 16 hours dark) with experimental temperatures (20° C, 12° C, 7° C). Body mass and the mass of food consumed by each animal were measured twice weekly for the duration of the experiment, and we note that – with very careful handling – it is possible to routinely weigh torpid animals without causing them to arouse from torpor (S3 Fig). Control groups that were housed under simulated summer conditions (~20° C, 16 hours light, 8 hours dark) maintained constant food consumption and body mass (Fig 5A). Body masses increased in the short photoperiod groups at 20 ° C, 12° C, and 7° C as the animals prepared for hibernation (Fig 5B-D). Using a threshold of a 25% percent body mass gain relative to starting condition as indicating the onset of preparation for hibernation, no control animals prepared while all of the experimental (short-photoperiod) animals prepared, except for one animal housed at 20° C (Table 2). Body masses reached a peak and fell once animals began to hibernate as indicated in Fig 5B-D. Food consumption remained constant in the control animals (Fig 5E), but increased in the experimental animals as they fattened for hibernation (Fig 5F-H). The food consumption of experimental animals was then greatly reduced as they spontaneously stopped eating and entered hibernation, as indicated in Fig 5F-H. We note that these animals do not store food in the hibernaculum in the wild, and would not normally emerge to forage during hibernation.

**Table 2.** Number of animals preparing for and entering hibernation by condition.

| Condition | Preparing | Hibernating | Total Animals |
| --- | --- | --- | --- |
| Control (20° C, 16L/8D) | 0 | 0 | 10 |
| Induction (20° C, 8L/16D) | 4 | 2 | 5 |
| Induction (12° C, 8L/16D) | 5 | 4 | 5 |
| Induction (7° C, 8L/16D) | 5 | 4 | 5 |

**Fig 5.**
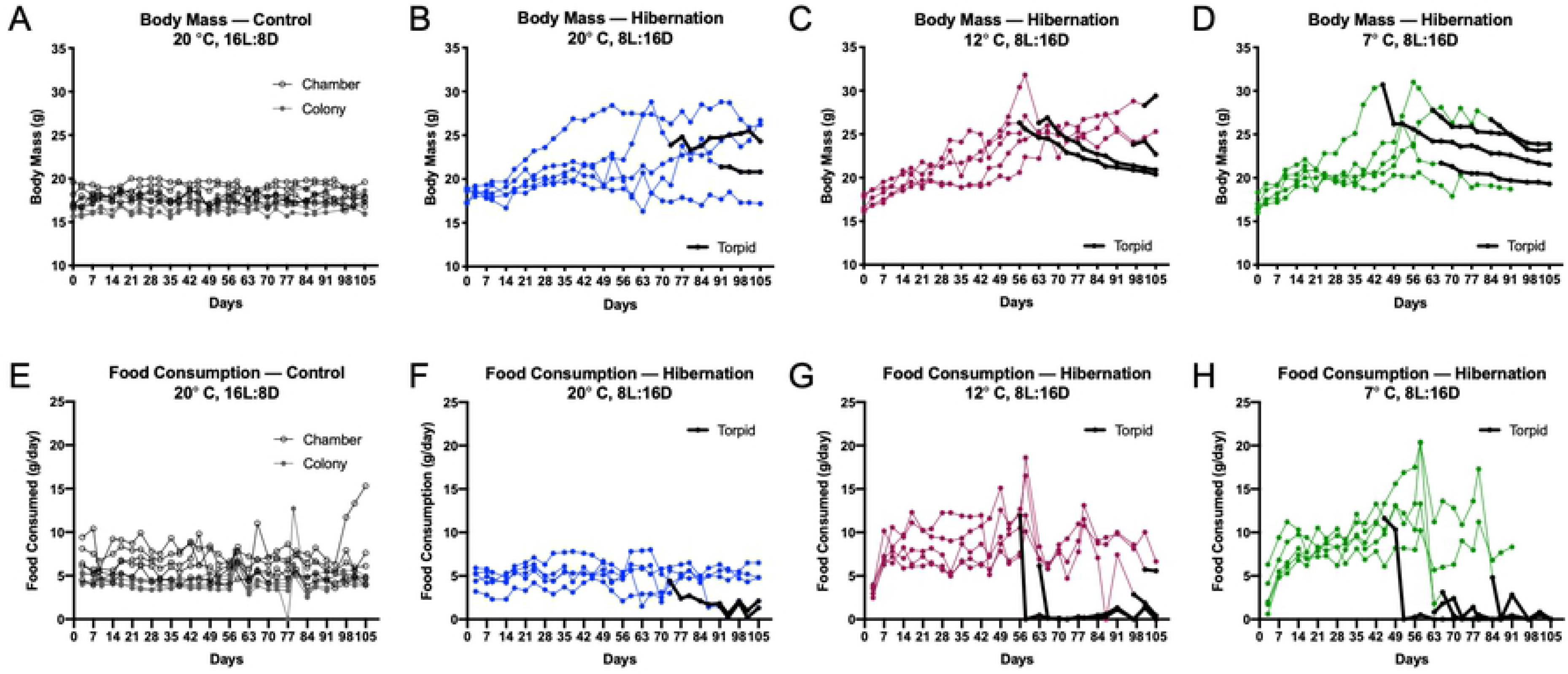
Short photoperiod and low temperature induce hibernation in captive-reared meadow jumping mice. Body mass (**A-D**) and average daily food consumption (**E-H**) are shown for control conditions (16L/8D, 20° C) and three hibernation induction conditions. Data are marked black when hibernation begins. (**A**) Body mass of control animals in an environmental chamber (open circles, n = 5), or colony housing room (closed circles, n = 5). (**B**) Body mass at 20° C and 8L/16D; 2 of 5 animals were found torpid. (**C**) Body mass at 12° C and 8L/16D; 4 of 5 animals were found torpid. (**D**) Body mass at 7° C and 8L/16D; 4 of 5 animals were found torpid. (**E**) Average daily food consumption of control animals in an environmental chamber (open circles, n = 5), or colony housing room (closed circles, n = 5). (**F**) Average daily food consumption at 20° C and 8L/16D; 2 of 5 animals were found torpid. (**G**) Average daily food consumption at 12° C and 8L/16D; 4 of 5 animals were found torpid. (**H**) Average daily food consumption at 7° C and 8L/16D; 4 of 5 animals were found torpid.

Additional analysis of the hibernation induction experiment provides insight into how temperature affects the food consumption and timing of hibernation and fattening in our animals. Short photoperiod appeared to be sufficient to drive fattening in all but one experimental animal (Table 2), while the number of animals found torpid in each experimental group was greater at lower temperatures (Table 2), confirming previous observations [21]. In the experimental animals, colder temperatures are additionally associated with a trend toward earlier onset of fattening and earlier hibernation (Fig 6A-B). Animals exposed to short photoperiod attained significantly greater maximum body mass at all temperatures compared to control animals (Fig 6C), but the number of days to maximum body mass did not differ by housing temperature (S4A Fig). In contrast, food consumption at maximum body weight showed an apparent dependence on temperature (S4B Fig). Experimental animals at 12° C and 7° C exhibited significantly greater food consumption at maximum body mass (Fig 6D), as might be expected due to the energetic demands of maintaining euthermic body temperature in the cold.

**Fig 6.**
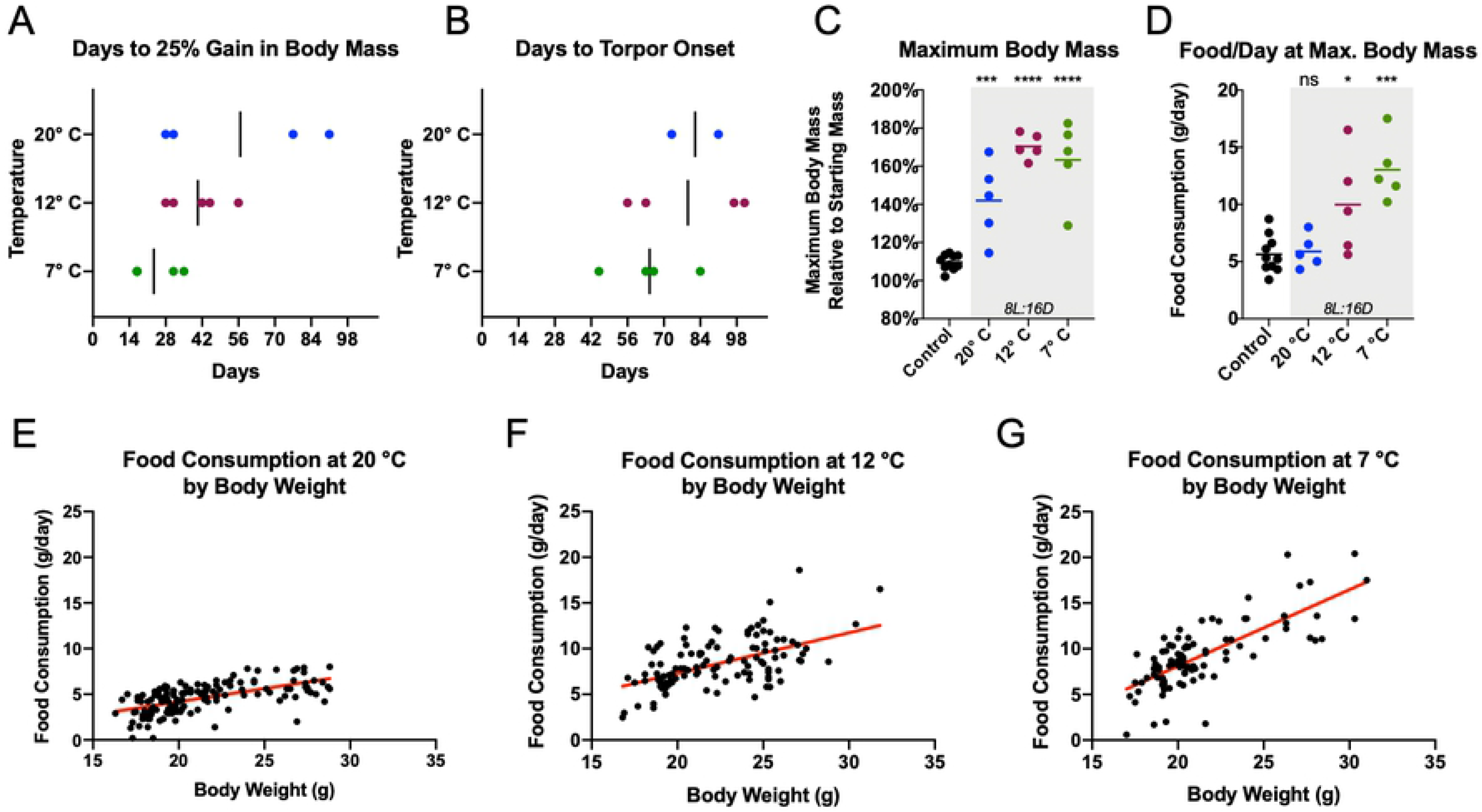
Ambient temperature affects pre-hibernation weight gain and food consumption in captive meadow jumping mice. Captive-reared meadow jumping mice in hibernation induction conditions (8L/16D and 20° C, 12° C, or 7° C) show trends toward earlier weight gain (**A**) and earlier onset of torpor (**B**) at lower temperatures. (**C**) Maximum attained body mass is significantly increased in all hibernation induction conditions (shaded gray) versus controls. Means differed significantly by ANOVA (F;30.92, p<0.0001) and asterisks indicate adjusted p values from Dunnett’s multiple comparisons test between control and each induction condition: 20° C (p;0.0007), 12° C (p<0.0001), and 7° C (p<0.0001). (**D**) Daily food consumption at maximum attained body mass is significantly increased at 7° C and 12° C, but not 20° C. Induction conditions are shaded gray. Means differed significantly by ANOVA (F;11.10, p;0.0001) and asterisks or ‘ns’ (not significant) indicate adjusted p values from Dunnett’s multiple comparisons test between control and each induction condition: 20° C (p;0.9964), 12° C (p;0.0166), and 7° C (p;0.0001) (**E-G**) The relationship between body weight and food consumption is shown at 20° C (panel **E**), 12° C (panel **F**), and 7° C (panel **G**). Red best fit lines increase in slope as temperature falls, illustrating the interaction between body mass and temperature with respect to food consumption.

To better understand the factors contributing to food consumption during preparation for hibernation, we analyzed the entire set of pre-torpor food consumption data for all groups using a multivariate model that included as factors body mass, photoperiod, temperature, housing location (environmental chamber vs. colony room), and litter for each animal, plus the interactions among the former three factors (S2 Table). The fit of the regression model was significant (p-value < 2.2 × 10^−16^, adjusted R^2^ = 0.6371). Factors with significant coefficients included body mass, temperature, the interaction between body mass and temperature, housing location, and some of the individual litters (S2 Table), suggesting that these factors play an important role in pre-torpor food consumption. Due to the significant interaction effect between body mass and temperature, these factors’ relationship to food consumption was non-linear. Plots of food consumption by body mass for all animals at each of the three temperatures (20° C, 12° C, and 7° C) allowed visualization of this phenomenon (Fig 6E-G). In these plots, the positive slope of the best-fit line increases as temperature falls, highlighting that 1) heavier animals exhibit greater food consumption, 2) the animals consume more food at colder temperatures, and 3) colder temperatures appear to have the greatest effect on food consumption by the heaviest animals. In other words, the colder the temperature, the greater the presumed energetic cost to the fattest animals. If this observation holds true in free-living meadow jumping mice, it would suggest that cold temperatures impose the greatest energetic cost on animals maintaining large fat stores at peak weight immediately before hibernation, and potentially help to explain the period of rapid weight loss immediately following initiation of hibernation observed by us and others (Fig 5C-D) [19, 40]. In addition to the factors of body mass and temperature, we note that the significant regression coefficients for litters of origin of the animals used in this study confirm the importance of genetic background on physiology among genetically diverse animals.

### Animal activity before and during hibernation

Other than a small set of 48-hour activity recordings previously reported by us [24], and one study of two western jumping mice (*Zapus princeps*) [48], we were not aware of any long-term activity monitoring data describing the circadian behavior of jumping mice. We therefore performed a long-term monitoring experiment on a group of five meadow jumping mice that were set to hibernate prior to a return to breeding. Following an initial acclimation period in the environmental chamber, the animals were housed at short photoperiod (8 hours light, 16 hours dark) and 7° C to induce hibernation. Food consumption was determined twice per week (as in the previous experiment), body mass was determined once per week to reduce potential disruption due to handling, and motion was recorded using a passive infrared motion detector mounted on the underside of the cage lid. The animals were provided a nest box for use as a hibernaculum, so the cage-top motion sensor only detected motion outside of the nest box. All five of the meadow jumping mice prepared for hibernation by gaining weight and 4 of the 5 animals were found torpid during the experiment (S5A-E Fig). An example animal with a robust hibernation response showed the characteristic pre-hibernation weight gain, then a hibernation interval marked by reductions in body mass and food consumption (Fig 7A) and in daily activity outside the nest box (Fig 7B). The animal that did not enter torpor maintained a steady body mass and food consumption during the study period (Fig 7C) and exhibited a relatively unchanging amount of daily motion outside of the nest box (Fig 7D). The activity data (Fig 7E-J and S5F-J Fig) collected from these animals confirmed the nocturnal behavior of *Zapus* spp. [24, 48] and revealed information about activity patterns before and during hibernation in captivity. While housed under simulated summer conditions (20° C, 16 hours light, 8 hours dark) during an initial acclimation period, the animals were typically active for the duration of the dark phase and remained in the nest box during the light phase (Fig 7E). Following the change to hibernation induction conditions (7° C, 8 hours light, 16 hours dark), and until the beginning of hibernation, the onset of activity typically aligned to the beginning of the dark phase, but the duration of activity did not expand to fill the complete 16-hour dark phase (Fig 7E-G). The start of hibernation coincided with cessation of most daily activity outside of the nest box, except for some sporadic periods of activity that were not aligned with the photoperiod (Fig 7H). Essentially no activity was recorded outside of the nest box for long periods of time during the middle of the recorded hibernation interval (Fig 7I), and late in hibernation we detected periods of activity outside the nest box that both did and did not align with the photoperiod (Fig 7J). Assuming that the mid-hibernation periods of activity outside of the nest box correspond to inter-bout arousals, these findings suggest that the meadow jumping mouse does not strictly align its arousals with the photoperiod occurring outside of the hibernaculum.

**Fig 7.**
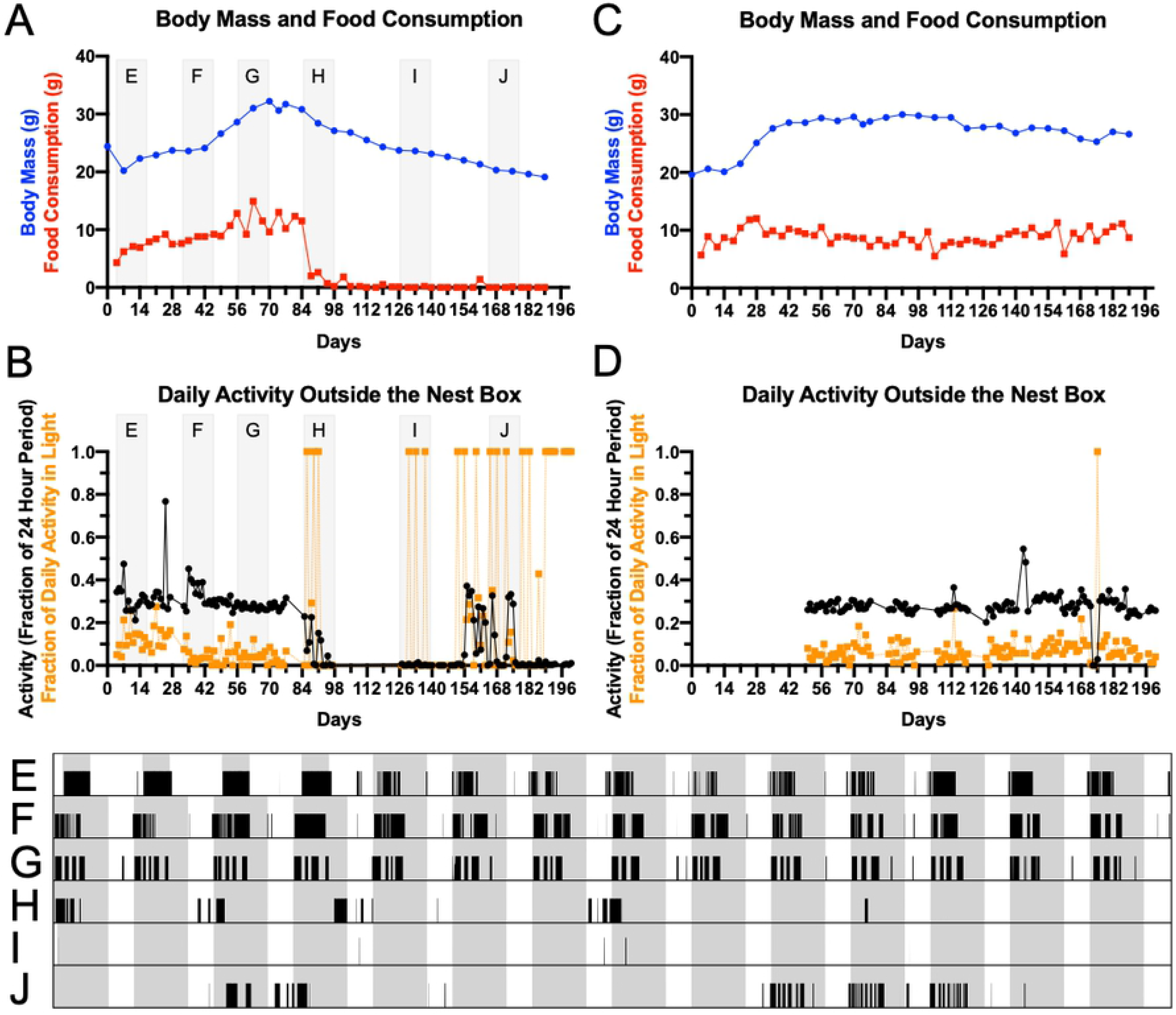
Circadian activity of captive meadow jumping mice before and during hibernation. Body mass (blue lines) and food consumption measurements (red lines) from animals that did (**A**) or did not (**B**) enter torpor at 8L/16D and 7° C. For the same animals that did (**C**) and did not (**D**) enter torpor, the total daily activity (fraction of 24 hour period active) is shown with black lines, and the fraction of total activity that occurred during the light phase is show with orange lines. The intervals shaded light gray in panels A and C correspond to the 14-day periods shown in actograms E-J. The dark phase is shaded gray and activity outside the nest box is represented as black bars. (**E)** First 14 days of activity. Hibernation induction conditions began after a 4 day acclimation period. **(F,G)** Before hibernation, daily activity begins at the onset of the dark phase. (**H**) At the start of hibernation, activity outside the nest box becomes sporadic and no longer aligns with photoperiod. (**I**) Mid-hibernation, essentially no activity occurs outside the nest box. (**J**) Late in hibernation, some activity occurs that both does and does not align with photoperiod.

To further understand the lengths of torpor bouts and arousals in the meadow jumping mouse, we used an uninterrupted recording of nest box temperature made during the hibernation of a female meadow jumping mouse (Fig 8A). The presence of a euthermic animal warms the nest box several degrees above ambient temperature, but a torpid animal does not produce sufficient heat to create a temperature differential; this phenomenon has been used previously to monitor the behavior of meadow jumping mice emerging from hibernation [49]. A torpor bout was deemed to begin when the nest box temperature differential (nest box temperature – chamber air temperature) first began to decline in the occupied nest box. The torpor bout ended, and the arousal was deemed to begin, when the temperature differential first began to increase above baseline. Using these measures, we quantified the lengths of 11 torpor bouts and 10 interbout arousals performed by the animal over a total hibernation interval of 90.6 days (Fig 8B-C). The torpor bout lengths increased for most of the hibernation interval from 2.1 days to a maximum of 12.8 days, then decreased prior to exit from hibernation (Fig 8B) in a pattern like that described previously [16]. Overall mean hibernation bout length was 7.7 days. Following an initial long (51.75 hour) arousal, arousal length remained relatively constant at a mean of 9.6 hours (range 7 – 12.25 hours) (Fig 8C). The lengths of torpor bouts and arousals in this animal are very similar to those of *Z. princeps* for the first 90 days of hibernation in that species [50]. We found that the timing of arousal onset relative to photoperiod drifted in a near-continuous fashion from mid-dark phase to mid-light phase over time (Fig 8D). Although the times of arousal onset in this animal did not occur “randomly” (i.e., were not uniformly distributed) throughout the day (p=0.017, Rayleigh test), data from additional animals and further study will be needed to understand the role, if any, daily external cues play in triggering arousals in the meadow jumping mouse.

**Fig 8.**
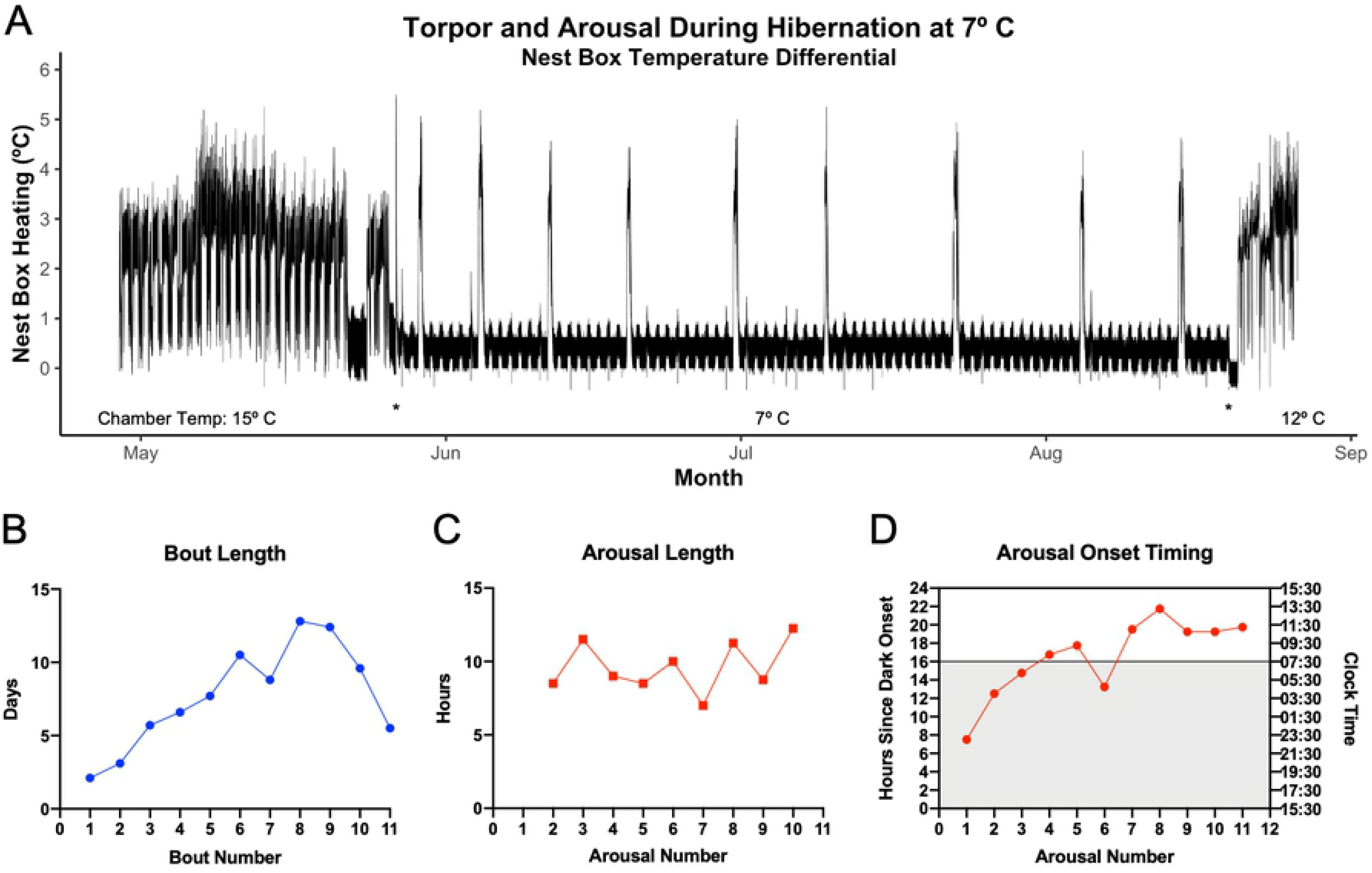
Duration of torpor bouts and arousals in a captive meadow jumping mouse. (**A**) Torpor and arousal as recorded by nest box temperature differential (° C) caused by body heat of the euthermic animal. Changes in the indicated ambient chamber temperatures are marked with asterisks. (**B**) Quantification of torpor bout length in days, as measured from onset of cooling to onset of warming of occupied hibernaculum. (**C**) Quantification of interbout arousal length in hours, measured from onset of warming to onset of cooling of occupied hibernaculum. (**D**) The timing of the beginning of each arousal. Time during the dark phase is shaded gray. Vertical axes show hours from onset of dark phase (left) and the corresponding clock time (right).

## Discussion

The male-skewed sex ratio and large number of single-sex litters weaned in our colony were unexpected given that field studies have found a balanced sex ratio of animals in the wild [20, 43, 51–53]. In laboratory mice, a diet high in saturated fat and low in carbohydrate results in a skew toward male offspring [54, 55]. If a similar mechanism is at work in our captive meadow jumping mice, perhaps the diet available in captivity, though nutritionally complete, is a poor match for the blend of carbohydrate, fat, and protein consumed by reproductively active animals in the wild. The varied reproductive strategies among mammals provide some additional possible explanations. An offspring sex ratio that varies over the active season – analogous to that seen in another hibernator, the big brown bat (*Eptesicus fuscus*) [56] – might offer an explanation if the environmental cues provided by our housing conditions interact with the seasonal biology of the meadow jumping mouse to favor production of male offspring. A skewed sex ratio alone, however, is not sufficient to explain the large numbers of all-male and all-female litters weaned in our colony. Although armadillos routinely produce single-sex litters consisting of monozygotic quadruplets [57, 58], the occurrence of monozygotic siblings is rare in other species with large litters, including the laboratory mouse [59], and the ability to experimentally produce single-sex litters in model organisms, including rodents, via genetic engineering is of current interest [60]. Among rodents, species of lemmings have evolved a chromosomal sex-determination system that allows some individuals to produce only female offspring [61]. Interestingly, a potentially complex system of sex determination involving XY males and XO females (as identified by karyotype) has been proposed for the western jumping mouse (*Zapus princeps*), but no karyotypic information is available for *Z. hudsonius* females [62]. The unexpected sex ratio and large number of single sex litters observed in our colony provide an opportunity to investigate factors controlling *Zapus* sex determination, the potential for the production of monozytogic siblings, or the possibility of selective removal of the minority sex from the litter by the female meadow jumping mouse.

Because meadow jumping mice are born throughout the summer, they are more dependent on environmental cues for timing of hibernation than are other species such as ground squirrels and marmots that exhibit strong circannual cycles controlling hibernation [63]. For example, Siberian chipmunks and golden-mantled ground squirrels exhibit free-running cycles of hibernation and activity with periods of typically slightly less than one year when held under constant conditions [64–66]. In contrast, when held under cold conditions of constant dark or short photoperiod, both meadow and western jumping mice appear to ‘skip’ the summer active phase and rapidly gain weight for repeated rounds of hibernation [21, 67], suggesting that they are capable of immediately responding to the environmental cues inducing hibernation without adhering to a strict circannual rhythm. The pre-hibernation fattening of the animals in our study that were captured late in the season but held at warm temperature and long photoperiod shows that the drive to hibernate is not reversed by external cues once it has begun, perhaps in a manner analogous to how photoentrainment in other rodent hibernators has little effect immediately prior to hibernation [63]. It thus appears that there exists in the meadow jumping mouse a physiological “hibernation program” that must run to completion once it has begun. While the mechanisms underlying the circannual rhythm in other hibernators are not known, we expect that the neurological mechanisms that translate short photoperiod to physiological changes in the meadow jumping mouse will share similarity with mechanisms of photoperiod control that are beginning to be revealed in seasonal breeders (e.g., Soay sheep, hamster) [68–70]. How the organism integrates the important seasonal cue of ambient temperature with photoperiod to allow appropriate timing of hibernation or other seasonal changes remains a mystery [71, 72], although important advances have been made in understanding temperature sensation in rodent hibernators [1].

The non-random timing of interbout arousals with respect to the photoperiod was somewhat surprising because we expected that the *Zapus* circadian clock would stop ticking at low body temperature as occurs in the hamster [73]; it stands to reason that, with protein translation in the torpid hibernator essentially halted [74], an oscillator based on the transcription and translation of clock genes would cease to oscillate. However, we note that a regular photoperiod was available as a potential zeitgeber during arousals, and we cannot rule out small daily fluctuations in temperature or other environmental inputs that have been shown to play a role in the timing of arousals in other hibernators including the mountain pygmy-possum [63, 75]. Future work using light-proof nest boxes or constant darkness during the hibernation interval (as would be experienced by the animal in a natural hibernaculum) will be needed to understand any circadian aspect of the timing of interbout arousals in the meadow jumping mouse.

The increase in food consumption exhibited by the pre-hibernation meadow jumping mouse at cold temperatures is similar in magnitude to the 2-3 fold increase in food consumption occurring in the thirteen-lined ground squirrel and reported for many hibernating species [76, 77]. Food consumption of our cold-housed meadow jumping mice seems to peak immediately prior to entry into hibernation, in a pattern that is unlike that seen in some larger rodent hibernators, where a slowing of metabolism that begins well before hibernation allows maximum weight gain to occur after the peak of food consumption [78]. We note that our meadow jumping mice fattened without large fold increases in food consumption when housed at 20° C, suggesting that cold temperature may impose a steep energetic penalty on this small hibernator during preparation for hibernation, a proposition that invites further investigation by measurement of energy expenditure via respirometry.

The short generation time of this species opens the door to genetic approaches that are otherwise difficult or impossible using hibernators with long generation times. The increasing facility of gene editing via nucleotide-directed endonucleases such as the CRISPR/Cas9 system may provide an opportunity to investigate hibernation phenotypes in *Zapus* using reverse genetics. Such an approach – for example, knocking out a gene of interest to observe the effect on a hibernation phenotype – would require a concerted effort to understand *Zapus* reproductive physiology and its compatibility with standard methods of gene editing that require manipulation of pre-implantation embryos, or the use of non-standard but potentially more feasible approaches to creating germline gene edits, such as microinjection and *in vivo* electroporation of adult testes or late-stage embryos with gene editing constructs [79, 80]. Once animals harboring a modified allele are available, the breeding strategies needed to generate study cohorts of appropriate genotypes are expected to be practicable. Beyond the promise of enabling hibernation genetics, the use of the meadow jumping mouse as a laboratory model of hibernation offers immediate opportunities for productive investigation of hibernation phenotypes.

## Acknowledgements

The authors thank Judith M. Chupasko, Mark D. Omura, Madeline D. Roorbach, L. Dwight Israelsen, Hilde R. Holcombe, James G. Fox, and Matthew G. Vander Heiden for assistance with this study.

## Supporting information

**S1 Fig. Simulated breeding outcomes and the histogram of litter size of all single-sex litters.** (**A**) Histogram of all possible gestation lengths as determined from the pairing and separation dates of 39 successful pairs of meadow jumping mice. The lower and upper bounds (shortest and longest possible gestation lengths) are overlaid in blue and pink, respectively. (**B**) Histogram showing number of litters by litter sex ratio, from 10 million sets of 32 litters that were simulated assuming an underlying sex ratio (fraction male) of 0.5. (**C**) Histogram showing number of litters by litter sex ratio, from 10 million sets of 32 litters that were simulated assuming an underlying sex ratio (fraction male) of 0.6615385, the ratio observed in our colony. (**D**) Plot showing the distribution of the number of single-sex litters obtained from 10 million sets of 32 litters simulated assuming an underlying sex ratio (fraction male) of 0.5. A red circle is placed at 19 single-sex litters, the number observed in our colony. (**E**) Plot showing the distribution of the number of single-sex litters obtained from 10 million sets of 32 litters simulated assuming an underlying sex ratio (fraction male) of 0.6615385, the ratio observed in our colony. A red circle is placed at 19 single-sex litters, the number observed in our colony. (**F**) A histogram of the sizes of 19 single-sex litters that occurred in our colony, by number of weaned pups per litter. The mean is 4 pups per litter.

**S2 Fig. Body mass of wild-caught and captive-reared meadow jumping mice.** (**A**) Each body mass measurements for each captive-reared animal from the time series shown in Fig 4A. *Zapus* born in captivity and held under simulated summer conditions (20 ° C, 16L/8D) maintained a relatively constant body mass for over one year. (**B**) Body mass over time for the animals captured during the second year of trapping and held at 20 ° C and 16L/8D photoperiod. For logistical reasons, data collection started some time after capture. These animals had fattened post-capture and then almost all of them spontaneously returned to a lean summer condition. (**C**) Body mass over time for the animals captured during the third year of trapping (black lines), or born to a captured female (red lines), and held at 16L/8D. For logistical reasons, data collection started some time after capture. Captured animals had fattened while captive-born animals maintained summer weight.

**S3 Fig. Example of weighing a torpid meadow jumping mouse directly on the scale.** This animal has rolled slightly onto its side on the laboratory scale; the head and feet are directly underneath the animal when hibernating in the hibernaculum. If handling must occur, accidental arousals can be minimized through rapid and gentle handling that does not uncurl the animal or turn it upside down.

**S4 Fig. Additional analysis of body mass and food consumption from a hibernation induction experiment.** (**A**) Days from start of hibernation induction to maximum attained body mass for each induction temperature. (**B)** Relationship between temperature and food consumed per day at maximum body mass during hibernation induction. Control animals are included at 20° C. Food consumption is increased at lower temperatures, as indicated by the negative slope of the best fit line.

**S5 Fig. Body mass, food consumption, and activity of all animals, related to Figure 7.** (**A-E)** Body mass and daily food consumption are shown for animals induced to hibernate in simulated winter conditions (7° C and 8L/16D). When applicable, black squares indicate the days the animal was found torpid. (**F-J**) Daily activity outside of the nest box as recorded by passive motion detector. Activity is represented as the fraction of each 24 hour day spent active by the animal (black lines and symbols), and the fraction of total activity occurring in the light phase is shown by orange lines and symbols. Before the hibernation interval, the animals typically spend less than half of the day active, with only a minor portion of total activity occurring during the light phase. Following entry into hibernation, a larger proportion of the reduced level of activity occurs during the light phase, consistent with disruption of the circadian rhythm.

**S1 Table. Pathogen testing results for animals captured each trapping season.**

**S2 Table. Regression results for pre-torpor food consumption.** Linear model of food consumption for each animal in the experiment shown in Fig 5.

